# Clock Modulation by Naringenin via RORα Suppresses Lipogenesis and Promotes Adipose Tissue Browning

**DOI:** 10.1101/2025.08.28.672947

**Authors:** Xuekai Xiong, Jemima Pangemanan, Tali Kiperman, Zuoming Sun, Antoni Paul, Vijay Yechoor, Ke Ma

## Abstract

The circadian clock orchestrates adipocyte development and lipid remodeling, with its disruption leading to the development of obesity and insulin resistance. Here we demonstrate that the flavonoid compound naringenin displays clock modulatory activity via RORα that suppresses adipocyte lipid storage while promoting browning. In adipogenic progenitors, naringenin activates RORα with induction of clock gene expression to promote circadian clock oscillation with protective effect against cytokine-induced dampening. The clock-enhancing properties of naringenin suppressed lipogenesis in mature adipocytes together with induction of browning characteristics. The inhibitory effect of naringenin on lipogenesis was dependent on clock modulation as it was abolished in RORα-deficient adipocytes. We further show that naringenin administration in vivo up-regulated RORα expression with clock gene induction together with browning of subcutaneous beige fat depot, resulting reduced fat mass and body weight. Naringenin treatment in vivo also lowered plasma glucose and free fatty acid levels, with markedly enhanced insulin signaling in adipose depots and skeletal muscle. Collectively, our findings uncover a new clock-activating mechanism of action in mediating the metabolic benefits of naringenin, suggesting its potential as a natural supplement for anti-obesity and metabolic disease interventions.

## Introduction

The circadian clock exerts rhythmic control in metabolic processes to coordinate systemic homeostasis in accordance with feeding and fasting nutrient oscillations (Bass & Takahashi, 2010; Takahashi, 2017). Misalignment of ∼24 hour endogenous circadian cycles with sleep-activity pattern or external nutrient cues, such as those caused by shiftwork or social “jet-lag”, disrupts metabolic homeostasis leading to the development of obesity and diabetes (Bass & Takahashi, 2010; Stenvers, Scheer, Schrauwen, la Fleur, & Kalsbeek, 2019). Maintenance of the temporal regulation of metabolic pathways may protect against clock disruption-induced metabolic disorders (Cederroth et al., 2019; Hatori et al., 2012; Sulli, Manoogian, Taub, & Panda, 2018).

Circadian clock-driven rhythmic oscillations of metabolic physiology are elicited by tightly coupled transcriptional and translational molecular feedback loops (Takahashi, 2017). Clock transcription activators, Bmal1 (Brain and Muscle Arnt-like 1) and CLOCK (Circadian Locomotor Output Cycles Kaput), form a heterodimer to activate the positive arm of the core molecular clock loop via induction of its direct target genes. Clock transcriptional repressors, including the Period and Cryptochrome genes, are transcription targets of CLOCK/Bmal1, whose activation initiates the negative feedback mechanism via repression of CLOCK/Bmal1 activity. Nearly all tissue and cell types possess cell-autonomous clocks that dictate daily oscillatory outputs, and the daily rhythmic regulations of distinct metabolic pathways collectively contribute to the appropriate timing of energy substrate metabolism (Bass & Takahashi, 2010).

In adipose tissue, adipogenic progenitors as well as mature adipocytes display robust cyclic rhythms (Otway, Frost, & Johnston, 2009; Zvonic et al., 2006). Key components of the molecular clock feedback loop are directly involved in modulating adipocyte development and function (Nam, Yechoor, & Ma, 2016). We previously reported that Bmal1, the essential clock activator, inhibits adipogenesis via the Wnt signaling pathway (Guo et al., 2012), while Rev-erbα, a direct target of Bmal1 and a transcriptional repressor of the re-enforcing arm of the clock regulatory loop, is required for brown adipogenesis (Nam, Chatterjee, et al., 2015). Interestingly, recent studies further implicated the coordinated circadian control in actin cytoskeleton-SRF/MRTF regulatory cascade that inhibits the development of thermogenic beige adipocytes (Xiong, Li, et al., 2023). Retinoid-related orphan nuclear receptors, RORα and RORγ, are known clock activators that induce clock gene transcription via a consensus RORE DNA-binding regulatory element (Jetten, 2009). Both RORs are strongly induced during adipogenic differentiation (Austin et al., 1998). Via direct interaction with a key early adipogenic transcription factor C/EBPβ, RORα blocks activities of PPARγ and C/EBPα to inhibit mature adipocyte terminal differentiation (Ohoka, Kato, Takahashi, Hayashi, & Sato, 2009). Interestingly, in the staggerer mice (Rorα^sg^) harboring a spontaneous deletion mutation of RORα, its deficiency resulted in thermogenic inductions of brown and beige adipocytes (Lau et al., 2015).

Disruption of clock can impair adipose tissue functions with pathological remodeling predisposing to the development of obesity (Karatsoreos, Bhagat, Bloss, Morrison, & McEwen, 2011; Kolbe, Leinweber, Brandenburger, & Oster, 2019). Chronic clock dysregulation by an experimental shiftwork regimen resulted in adipose tissue expansion accompanied by inflammation and fibrosis with impaired systemic insulin sensitivity (Xiong et al., 2021). Conversely, re-enforcing clock function, such as via time-restricted feeding in high fat diet-fed animal model, protects against the development of obesity with insulin-sensitizing actions (Hatori et al., 2012). Despite accumulating studies indicating that maintaining or modulating clock activity may yield metabolic benefits, clock-targeting interventions for obesity and diabetes remain scarce. Rev-erbα activation by synthetic agonists demonstrated anti-obesity and anti-lipidemic effects (Solt et al., 2012). We recently found that Nobiletin, a citrus flavonoid known to enhance clock cycling amplitude, displays direct anti-adipogenic effect in a clock-dependent manner (He et al., 2016), demonstrating the potential for discovery of clock-modulatory molecules as therapeutic avenues against metabolic diseases (Chen, Yoo, & Takahashi, 2018).

Naringenin (NAR) is one of the polyphenolic flavonoid compounds highly abundant in citrus fruits (Tsao, 2010). Epidemiological studies revealed that intake of flavonoid-rich food is associated with metabolically beneficial effects on cardiovascular disease, dyslipidemia and Type II diabetes (Cassidy et al., 2012; Hooper et al., 2008; Wedick et al., 2012). Our recent findings revealed that Nobiletin, a clock amplitude-enhancing flavonoid via RORα/γ activation, (Assini, Mulvihill, & Huff, 2013; Nohara et al., 2019), exerts direct inhibition of adipogenesis that reduces adipose tissue expansion in vivo (He et al., 2016). As a flavonoid compound with related structural scaffold with Nobiletin, Naringenin is known to display beneficial metabolic properties (Assini et al., 2013), though its precise mechanisms of action in adipocyte biology remains to be explored. Based on the role of clock in adipocyte development and the closely related structural scaffold between NAR and Nobiletin (Guo et al., 2012; Nam, Chatterjee, et al., 2015; Xiong, Li, et al., 2023; Zhang, Sinha, Bahrami-Nejad, & Teruel, 2022), we tested whether Nar possesses clock-modulatory activities that may influence adipocyte function. Our findings identified NAR as a new ROR-activating natural compound with clock-dependent actions in adipocyte lipid synthesis and thermogenic browning that contribute to its anti-obesity efficacy in vivo.

## Materials & Methods

### Animals and drug delivery

Mice were maintained in the City of Hope vivarium under a constant 12:12 light dark cycle, with lights on at 6:00 AM. All animal experiments were approved by the Institutional Animal Care & Use Committee (IACUC) of City of Hope and carried out in concordance with the IACUC approval. C57BL/6J mice were purchased from Jackson Laboratory and used for experiments following 2 weeks or longer of acclimation at 10 or 48 weeks of age. Naringenin was purchased from Santa Cruz (CAS 480-41-1). Mice were given an intraperitoneal injection daily with NAR (50 mg/kg) or vehicle (DMSO) for 7 days. Floxed RORα mouse line was a gift from Dan Littman at NYU (Hall et al., 2022). Adipocyte-specific RORα-null mice were generated by crossing the floxed RORα allele with Adipoq-Cre transgenic mice (Jackson Lab Strain 028020).

### Cell culture

3T3-L1 preadipocytes (RRID:CVCL_0123) were obtained from ATCC, and maintained in DMEM with 10% fetal bovine serum supplemented with Penicillin-Streptomycin-Glutamine, as previously described (Liu, Xiong, Nam, Yechoor, & Ma, 2020; Nam, Guo, et al., 2015). Cells were seeded at 1×10^6^ density on 6-well plates and treated with naringenin (5uM and 10 μM) or vehicle (DMSO) for 24 hours prior to protein extraction.

### Primary preadipocyte isolation and treatment

The stromal vascular fraction containing preadipocytes were isolated from subcutaneous fat pads, as described (Xiong, Li, et al., 2023). Briefly, fat pads were dissected into ∼0.5 mm pieces and digested using 0.1% collagenase Type 1 with 0.8% BSA at 37^0^C in a horizontal shaker for 60 minutes. Homogenates were filtered through a Nylon mesh and centrifuged to obtain the pellet containing the stromal vascular fraction with preadipocytes. Preadipocytes were cultured in F12/DMEM supplemented with bFGF (2.5 ng/ml) and expanded for two passages before being seeded for experiments in 6-well plates 1×10^6^ density. Preadipocytes were treated with naringenin (5uM and 10uM) or vehicle (DMSO) for 8 hours, before addition of cytokine cocktail (IL1β-10 ng/ml, IFNγ-100 ng/ml, and TNFα-25ng/ml) for an additional 16 hours prior to protein extraction.

### Primary preadipocyte differentiation

Primary preadipocytes isolated from mice subcutaneous fat pads were subjected to differentiation in 6-well plates at 90% confluency. Adipogenic differentiation was induced for 2 days in medium containing 10% FBS, 1.6 μM insulin, 1 μM dexamethasone, 0.5 mM IBMX, 0.5 µM rosiglitazone before switching to maintenance medium for 4 days with insulin only. Naringenin (5uM and 10uM) or vehicle (DMSO) were added for the entire differentiation time course at beginning of adipogenic induction. Beige adipocyte differentiation was induced for 2 days using induction medium (10% FBS, 1.6 μM insulin, 1 μM dexamethasone, 0.5 mM IBMX, 0.5 μM Rosiglitazone, and 1 nM triiodothyronine) before switching to maintenance medium (10% FBS with insulin, Rosi, and T3 only) for 4 days. Naringenin (5uM and 10uM) or vehicle (DMSO) were added at 4 day-differentiated beige adipocytes for 2 days prior to staining and protein extraction.

### Oil-red-O, Bodipy and MitoTracker staining

These staining for neutral lipids and mitochondria in differentiated adipocytes were performed as previously described (Nam, Guo, et al., 2015). Briefly, for oil-red-O staining, cells were fixed using 10% formalin and incubated in 0.5% oil-red-O solution for 1 hour. Bodipy 493/503 (RRID: AB_2536193) was used at 1mg/L together with DAPI for 15 minutes, following 4% paraformaldehyde fixation and permeabilization with 0.2% Triton X-100. MitoTracker Deep Red FM (ThermoFisher) was applied at 100 nM to differentiated adipocytes and incubated for 30 min prior to fixation. Bodipy and MitoTracker fluorescence intensity were measured using Image J for quantitative analysis of (n=3/group).

### Continuous Bioluminescence monitoring of Per2::dLuc luciferase reporter in isolated primary adipocytes

Primary preadipocytes isolated from mice containing a *Per2::dLuc* luciferase reporter transgene were seeded at 4×10^5^ density on 24 well plates. Preadipocytes were treated with naringenin (2-10 µM) or vehicle (DMSO) in the presence or absence of a cytokine cocktail (10 ng/ml IL1β, 100 ng/ml IFNγ, and 25ng/ml TNFα). Real-time luciferase activities were recorded for 7 days using LumiCycle 96 (ActiMetrics), as previously described (Xiong et al., 2022). Briefly, cells were used at 90% confluence following overnight culture with explant mediµM luciferase recording media. Explant medium contains 50% 2xDMEM buffer stock, 10% FBS (Cytiva), 1% PSG, pH7 1M HEPES, 7.5% Sodium Bicarbonate, sodiµM hydroxide (100 mM) and XenoLight D-Luciferin bioluminescent substrate (100 mM). Raw and subtracted results of real-time bioluminescence recording data for 6 days were exported, and data was calculated as luminescence counts per second. LumiCycle Analysis Program (ActiMetrics) was used to determine clock oscillation period, length amplitude and phase. Briefly, raw data following the first cycle from day 2 to day 5 were fitted to a linear baseline, and the baseline-subtracted data (polynomial number = 1) were fitted to a sine wave, from which period length and goodness of fit and damping constant were determined. For samples that showed persistent rhythms, goodness-of-fit of >80% was usually achieved.

### ROR-responsive luciferase reporter assay

RORE-containing luciferase reporter containing three RORE bindings sites RORE(3) TK-luc was a gift from Zuoming Sun lab (Kane & Means, 2000; Medvedev, Yan, Hirose, Giguere, & Jetten, 1996). 293T cells were seeded and grown overnight to ∼70% confluency prior to plasmid transfection. Cells were transfected using PolyJet reagent (SignaGen Laboratories) with expression plasmids, including RORE-luciferase reporter (50 ng/well), Renilla (20 ng/well), and RORα (20 ng/well) following the manufacturer’s protocol. 24 hours following transfection, cells were treated with NAR or SR1078 at indicated concentrations overnight to induce luciferase activity. Luciferase activity was assayed using Dual-Luciferase Reporter Assay Kit (Promega) in 96-well black plates. Firefly luciferase reporter luminescence was measured on microplate reader (TECAN infinite M200pro) and normalized to control Renilla luciferase activity, as previously described (Guo et al., 2012). The mean and standard deviation values of four repeats were calculated for each well and graphed.

### Immunoblot analysis

Protein was extracted using lysis buffer (3% NaCl, 5% Tris-HCl, 10% glycerol, 0.5% Triton X-100 in sterile Milli Q water) containing protease inhibitors. 20-40 µg of total protein was resolved on 10% SDS-PAGE gels followed by immunoblotting on PVDF membranes (Bio-rad). Membranes were developed by chemiluminescence (SuperSignal West Pico, Pierce Biotechnology) and signals were obtained via a chemiluminescence imager (Amersham Imager 680, GE Biosciences). Antibodies used were listed in Supplemental Table 1.

### RNA extraction and RT-qPCR analysis

Trizol (Invitrogen) and PureLink RNA Mini Kit (Invitrogen) were used to isolate total RNA from snap-frozen tissues or cells, respectively. cDNA was generated using Revert Aid RT kit (ThermoFisher) and quantitative PCR was performed using SYBR Green Master Mix (Thermo Fisher) in triplicates on ViiA 7 Real-Time PCR System (Applied Biosystems). Relative expression levels were determined using the comparative Ct method with normalization to 36B4 as internal control. PCR primers sequence were listed in Supplemental Table 2.

### Hematoxylin and eosin histology

Adipose, skeletal muscle or liver tissues were fixed in 10% neutral-buffered formalin for 72 hours prior to embedding. 10 μm paraffin sections were processed for hematoxylin and eosin staining.

### Plasma metabolite analysis

Plasma levels of glucose (Thermo Scientific), triglyceride (Teco Diagnostics) and free fatty acids (BioAssay Systems) were measured using 5-10μl of plasma samples with respective commercial kits, according to manufacturer’s protocols.

### Statistical analysis

Data are presented as mean ± SE as indicated. Each in vitro experiment was repeated at a minimum of three times to validate the results. Sample sizes are indicated for each experiment in figure legends. Statistical analysis was performed using GraphPad Prism10 software. A minimum of three biological replicates were used to perform statistical analysis. Two-tailed Student’s t-test or one-way ANOVA with Tukey’s post-hoc analysis for multiple comparisons were performed as indicated. P<0.05 was considered statistically significant.

## Results

### The citrus flavonoid Naringenin activates RORα and promotes clock oscillation in adipogenic progenitors

Naringenin is a hydrophilic flavonoid compound with a core scaffold resembling that of Nobiletin (Fig. 1A). Based the known function of Nobiletin as a RORα-targeting clock modulator that enhances clock cycling amplitude (He et al., 2016), we explored potential clock-modulatory activities of NAR and tested whether it can function as a RORα ligand. Using a RORE-containing luciferase reporter construct (Kane & Means, 2000), we examined the activity of NAR together with a RORα synthetic agonist SR1078 as a positive control (Wang et al., 2010). As expected, the transfection of RORα is sufficient to induce RORE-Luc activity by ∼15 fold over basal empty vector control (Fig. 1B). Nar treatment at 5µM was able to further activate RORE-driven luciferase reporter by to ∼23 fold over basal level. Interestingly, the activity of Nar at 5µM was comparable to that of the activation by SR1078 at 1µM. Thus, the natural flavonoid NAR displayed bona fide, albeit lower, RORα agonist activity. In 3T3-L1 preadipocytes, NAR led to a robust dose-dependent induction of RORα protein (Fig. 1C). NAR similarly induced Bmal1 expression, potentially due to RORα activation of Bmal1 as a direct target in the core clock feedback loop (Takahashi, 2017). These findings thus implicated NAR as a circadian clock modulator via RORα activation. We next determined the direct modulatory effect of NAR on clock oscillation via continuous real-time monitoring of *Per2::dLuc* luciferase reporter activity using primary preadipocytes isolated from *Per2::dLuc* transgenic mice (Fig. 1D-1F). Chronic low-grade inflammation in adipose tissue is a key feature of human obesity (Saltiel & Olefsky, 2017; Wu & Ballantyne, 2020). Based on the known anti-inflammatory activity of NAR, we also examined its potential protective effects against a cytokine cocktail (CC) challenge mimicking an inflammatory milieu (Crewe, An, & Scherer, 2017; Guzik, Skiba, Touyz, & Harrison, 2017). NAR treatment induced a dose-dependent increase of clock cycling amplitude at 2 to 5 µM (Fig. 1D & 1E), although it did not alter period length (Fig. 1F). In contrast, cytokine treatment led to marked dampening of clock amplitude and reduction of period length. Notably, NAR partially rescued CC-induced reduction of clock amplitude in these preadipocytes (Fig. 1E). In primary preadipocytes, NAR was also able to induce RORα and Bmal1 protein expression, more robustly at 10 µM than 5 µM (Fig. 1G). Furthermore, NAR conferred resistance to CC-induced attenuation of these clock regulators, in line with its ability to rescue clock oscillatory amplitude under this condition. Together, these findings identify NAR as a novel RORα-activating flavonoid that promotes clock oscillation.

**Figure 1.**
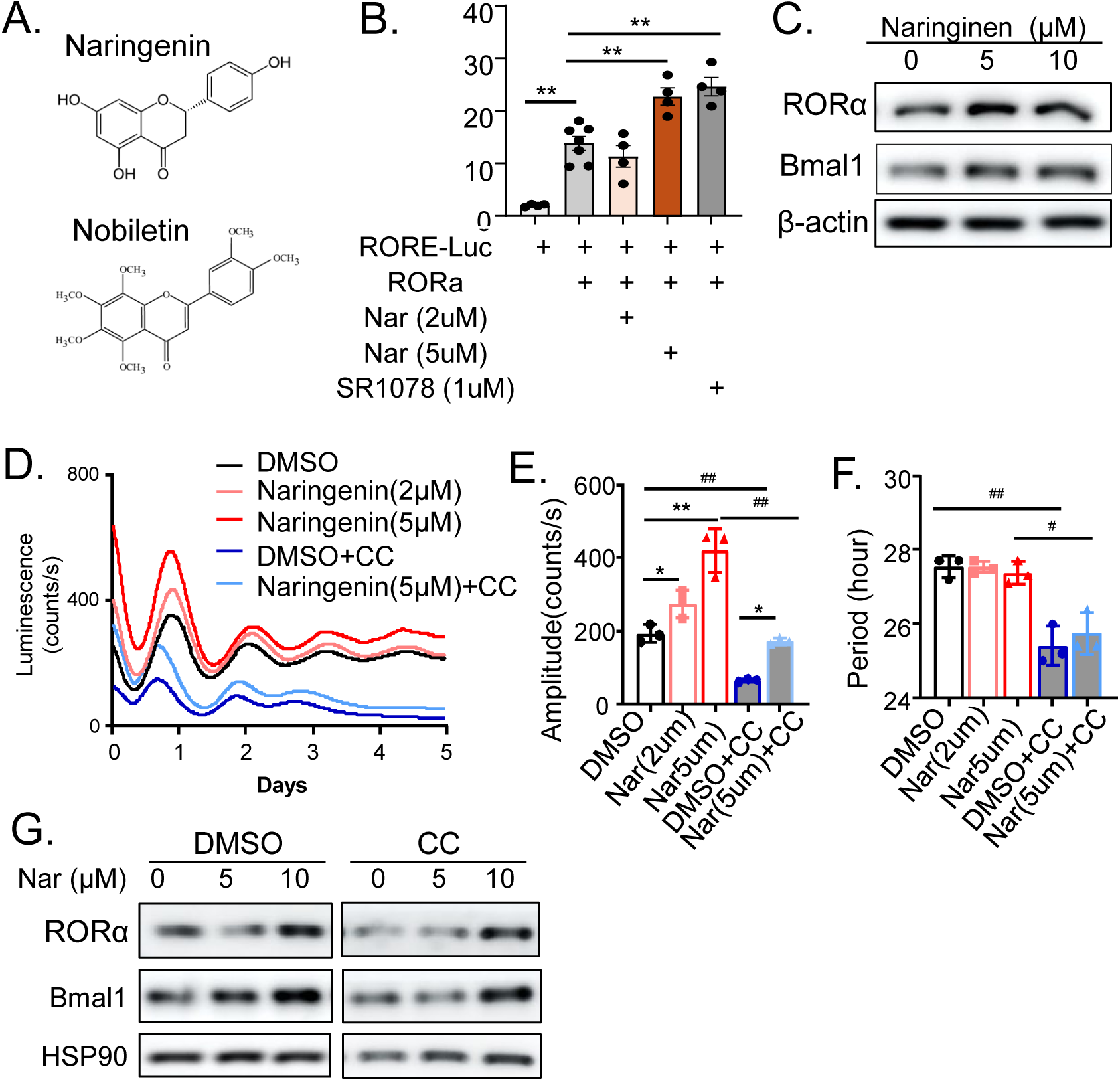
Naringenin promotes clock oscillation and protects against inflammatory cytokines. (A) Chemical scaffold of Naringenin in comparison with Nobiletin. (B) Effect of Naringenin and RORα agonist SR1078 on RORE-responsive luciferase reporter activity. N=4/groups. *: P≤0.01 by ANOVA. (C) Immunoblot analysis of Naringenin effect on induction of clock genes in 3T3-L1 preadipocytes at indicated concentrations. (D-F) Effect of Naringenin on clock modulation monitored by continuous real-time chemiluminescence recordings of preadipocytes containing a *Per2-dLuc* reporter for 5 days. Average tracing of luciferase reporter at indicated Naringenin concentrations with or without cytokine cocktail (CC: IL-1β, IL6 and TNFα) (D), and quantitative analysis of clock oscillation amplitude (E) and period length (F). N=4/group. Results are presented as Mean ± SE of n=3 replicates. *, **: p<0.05 and 0.01 by one-way ANOVA with Tukey’s post-hoc analysis. (G) Immunoblot analysis of Naringenin effect on clock genes in primary preadipocytes at indicated concentrations, treated with or without CC.

### Naringenin suppresses lipogenesis and induces browning of mature adipocytes

Our previous study revealed that Nobiletin, a known RORα agonist, displayed clock-dependent inhibitory effect on adipogenesis (Xiong, Kiperman, et al., 2023). We postulated that due to its clock-enhancing properties through RORα, NAR, may impact adipocyte development. We thus treated primary preadipocytes undergoing a standard white adipocyte differentiation regimen with NAR at increasing concentrations, and assessed their effects on adipocyte development through analyses of phase-contrast morphology, along with oil-red-O and Bodipy staining to determine lipid accumulation (Fig. 2A). Interestingly, there were no significant effects of 5 or 10 µM NAR treatment on adipocyte maturation under this differentiation condition, as indicated by comparable levels of lipid accumulation as compared to vehicle controls that was confirmed by quantification of Bodipy staining (Fig. 2B). We next tested the effects of NAR on mature adipocytes by treating them following differentiation and determined its potential impact on the inducible thermogenic characteristics of beige adipocytes, namely browning. Primary preadipocytes were induced to differentiate into mature beige adipocytes with addition of T3 in the adipogenic cocktail before NAR were added at indicated concentrations for 2 days. Surprisingly, under this beige adipogenic condition, NAR displayed a marked effect on suppressing lipid storage in differentiated adipocytes, as shown by the significant reduction of Bodipy-stained lipids at 10 µM (Fig. 2C & 2D). Furthermore, staining for active mitochondria using MitoTracker revealed significantly elevated mitochondrial mass by both 5 and 10 µM of NAR treatment (Fig. 2C & 2D), suggesting a browning effect with elevated mitochondrial mass/activity under this condition. Consistent with the observed reduction in lipids by NAR, analysis of protein expression revealed suppression of genes involved in lipogenesis by 10 µM of NAR, including fatty acid binding protein 4 (FABP4) and fatty acid synthetase (FASN), whereas the adipogenic factor C/EBPβ was not altered (Fig. 2E). In contrast, lipolytic enzymes involved in mobilization of lipids from adipocytes, including adipose triglyceride lipase (ATGL), hormone-sensitive lipase (HSL) and their phosphorylation, were induced by NAR (Fig. 2F). The observed NAR effect on inducing browning was further corroborated by the up-regulation of a key effector of thermogenesis, the uncoupling protein-1 (UCP-1) protein, and elevated expression of mitochondrial markers, including succinate dehydrogenase subunit B (SDHB) and ATP5a while peroxisome proliferator-activated receptor gamma coactivator-1 alpha (PGC-1α) was also increased (Fig. 2G). Thus, the attenuated lipid storage by NAR in mature adipocytes could be due to suppression of lipogenesis in parallel with lipolytic activation, while induction of browning may further contribute to the depletion of lipids.

**Figure 2.**
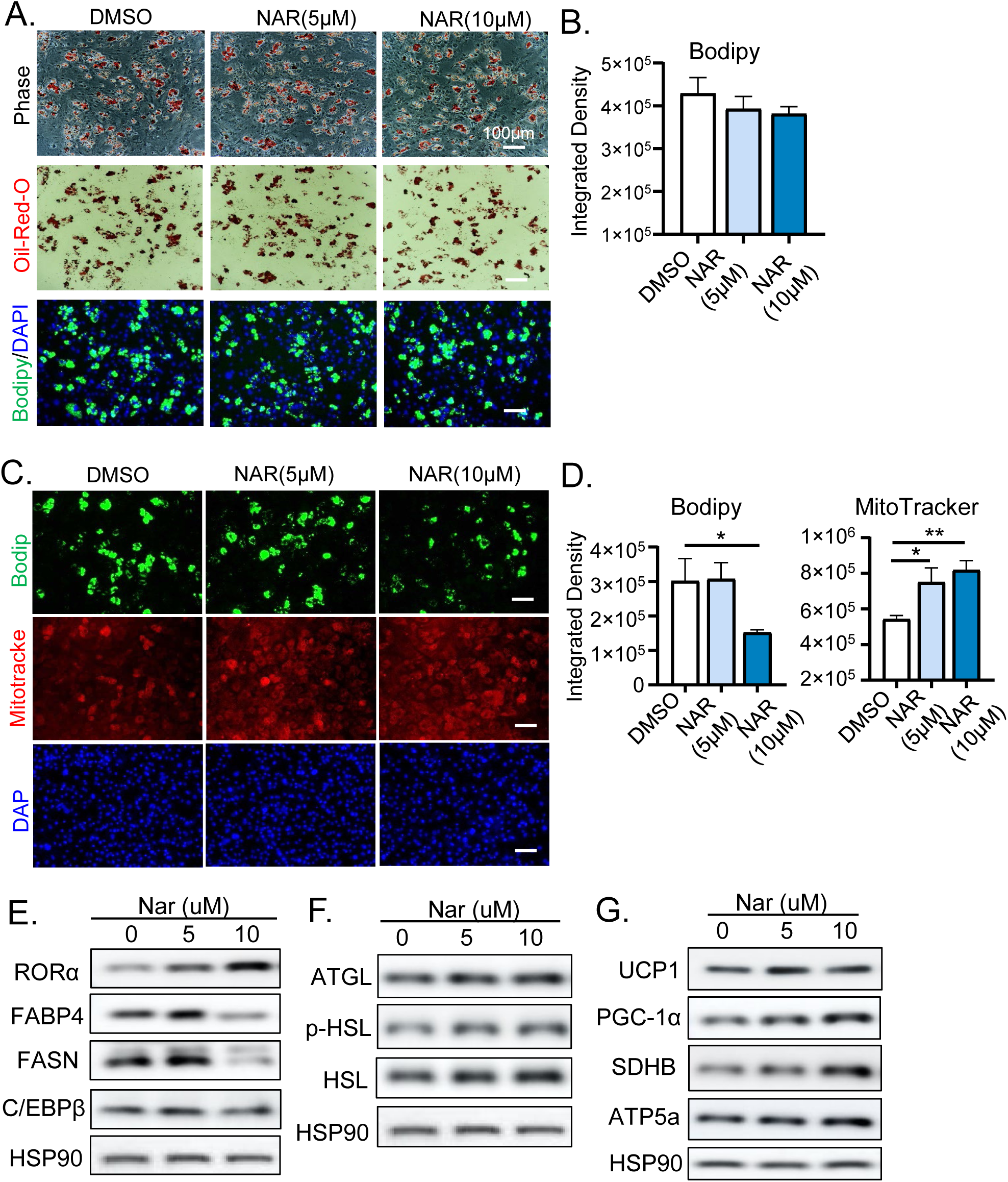
Effect of Naringinen on adipogenic differentiation and browning of mature adipocyte. (A, B) Representative images of phase-contrast, oil-red-O and Bodipy staining of primary preadipocytes following 6 days of differentiation, in the presence of Naringenin (NAR) at indicated concentrations (A), with quantitative analysis of Bodipy staining (B). (C) Representative images of Bodipy and MitoTracker staining of 6-day differentiated beige adipocytes treated with 5 &10μM Naringenin during last 2 days of differentiation (day 4 to 6), with quantitative analysis (D). (E-G) Immunoblot analysis of lipogenic (E), lipolytic (F) and thermogenic gene markers (G) following Naringenin treatment of differentiated beige preadipocytes for 48 hours. Each lane represents a pooled sample of three independent experiments.

Given that NAR stimulated RORα activity and was sufficient to induce its expression, we next determined whether its effect on suppressing lipid accumulation or browning of beige adipocytes is dependent on RORα. Using primary preadipocytes isolated from mice with adipocyte-specific RORα ablation, through a cross of adiponectin-Cre transgenics with floxed RORα mice (Hall et al., 2022), we tested the RORα-dependency of NAR activity in differentiated beige adipocytes. Floxed control or RORα-deficient preadipocytes were subjected to beige differentiation followed by NAR treatment for 48 hours. Similarly as seen in wild-type beige adipocytes, NAR significantly inhibited lipid accumulation in floxed control adipocytes (Fig. 3A & 3C). The effect on inhibiting lipid storage by NAR was completely abolished in RORα-deficient adipocytes, suggesting that this activity is dependent on RORα (Fig. 3B &3C). NAR also exhibited a trend toward enhancing mitochondrial activity as shown by MitoTracker staining, though not significant as compared with controls based on quantitative analysis (Fig. 3A & 3D). These findings revealed the direct modulatory effect of NAR in mature adipocytes without altering their differentiation, with its inhibition on lipid storage dependent on RORα regulation.

**Figure 3.**
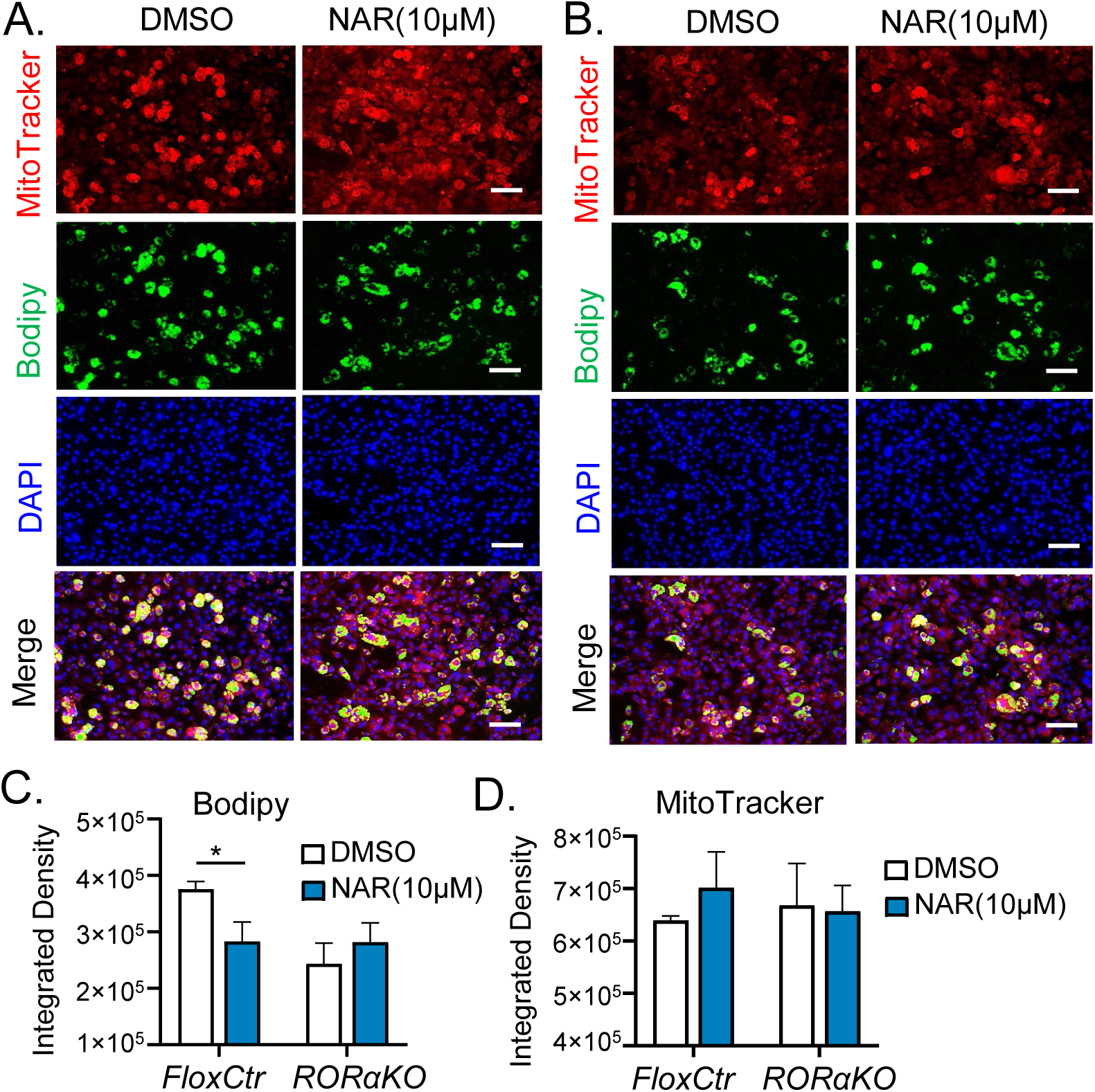
RORα-dependency of Naringinen effect on suppressing lipid storage in adipocytes. (A, B) Representative images of Bodipy and MitoTracker staining of Flox and RORα-deficient beige adipocytes following Naringenin treatment at day 6. Primary preadipocytes isolated from inguinal subcutaneous depots of control Floxed RORα mice (FloxCtr, A), and adipocyte-specific RORα-deficient mice (RORαKO, B) were differentiated into beige adipocytes for 6 days with Naringenin (10 μM) treatment during day 4 to 6. Scale bar: 100 μm. (C & D) Quantitative analysis of Bodipy (C) and MitoTracker staining (D) of Flox and RORα-deficient beige adipocytes with or without Naringenin treatment. n=3/group.

### In vivo efficacy of Naringenin on reducing fat mass and improving glucose metabolism

Based on NAR suppression of lipid accumulation with induction of browning in mature adipocytes, we next determined its effect on distinct fat depots in normal wild-type C57/BL6 mice. Intraperitoneal delivery of NAR for 7 days led to significantly reduced body weight in treated cohorts, while the body weight of vehicle-treated controls remained comparable with before treatment (Fig. 4A). Examination of inguinal white adipose tissue (iWAT) beige fat depot revealed significantly lower mass than that of controls (Fig. 4B), while visceral epididymal white adipose tissue (eWAT) was comparable (Fig. 4C). Interestingly, the classic thermogenic fat interscapular brown adipose tissue (BAT) weight was also reduced, suggesting predominant effect of NAR on thermogenic fat depots (Fig. 4D). In addition, liver weight was also reduced by NAR treatment (Fig. 4E), suggesting potentially attenuated fat accumulation in the liver. In contrast, heart and muscle weights, such as the gastrocnemius muscle (GN), were similar to vehicle-treated control mice (Fig. 4F & 4G), suggesting that loss of fat by NAR treatment likely account for majority of the observed reduction in total body weight. NAR also resulted in significant beneficial effects on systemic metabolic regulations. This included lower plasma blood glucose level (Fig. 4H) and circulating free fatty acid (Fig. 4I), although plasma triglyceride level was similar to controls (Fig. 4J). The glucose-lowering effect of NAR suggests potential improvement in insulin sensitivity. We thus examined insulin signaling and found that Akt phosphorylation in both inguinal beige fat and interscapular brown fat were markedly enhanced (Fig. 4K & 4L). In addition, total Akt protein level in the iWAT was induced by NAR treatment but not in the BAT. Similarly elevated Akt phosphorylation was observed in skeletal muscle (Fig. 4M), suggesting systemic improvement in insulin sensitivity as a result of NAR treatment that may mediate enhanced glucose utilization.

**Figure 4.**
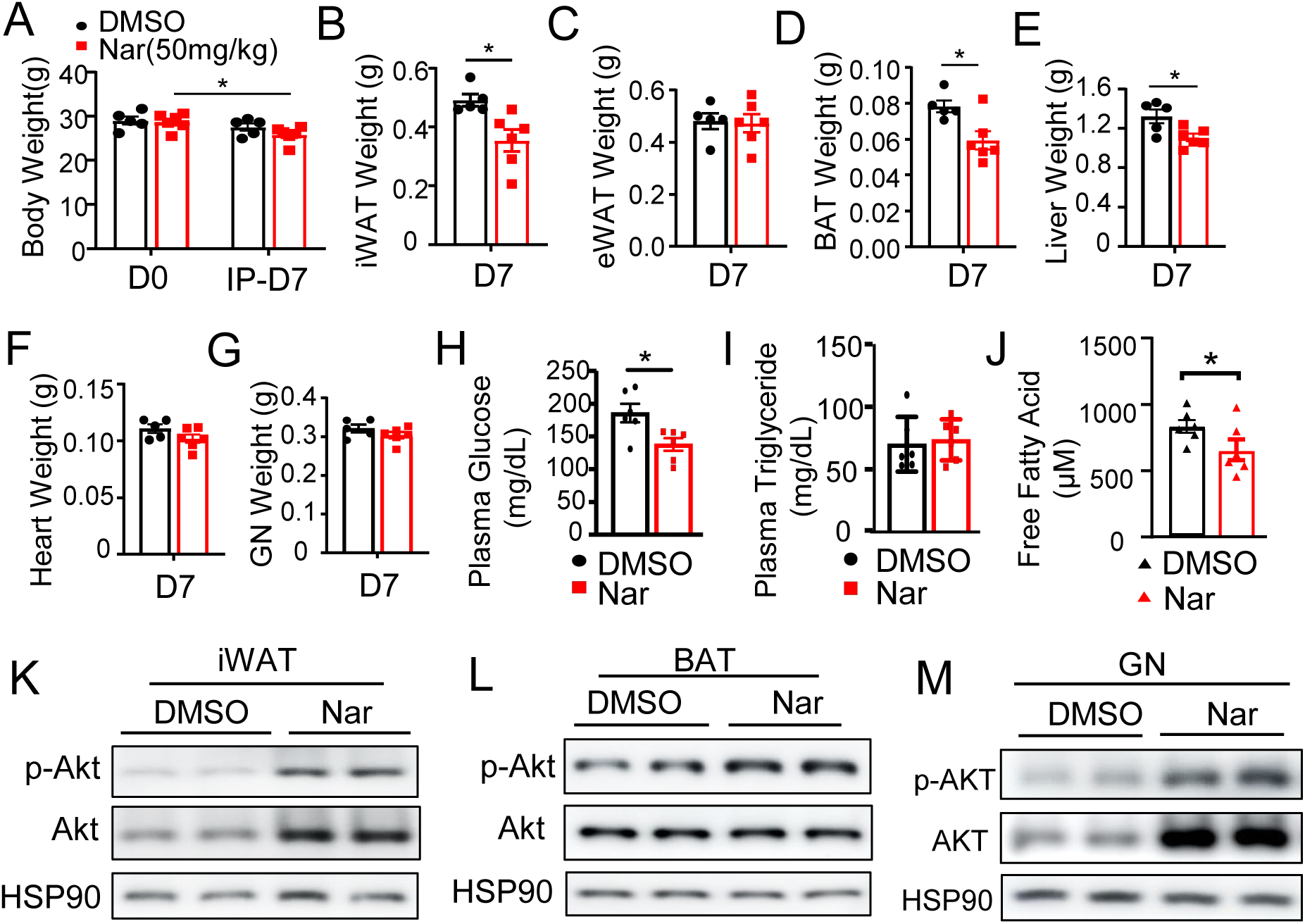
In vivo effect of Naringenin on reducing fat mass and promoting insulin signaling. (A) Effect of Naringenin treatment on body weight via daily intraperitoneal injection (50mg/kg) for 7 days. Male C57/BL6 mice at 12 weeks of age were used (n=6/group). (B-G) Analysis of tissue weight following 7 days of Naringenin administration, including inguinal white adipose tissue (iWAT, B), visceral epidydimal white adipose tissue (eWAT, C), interscapular brown adipose tissue (BAT, D), liver (E), heart (F), and gastrocnemius muscle (GN, G). (H-J) Systemic metabolic effect of Naringenin on plasma glucose (H), triglyceride (I) and free fatty acid levels (J) following in vivo administration. *: P≤0.01 by Student’s t test. (K-M) Immunoblot analysis of Naringenin effect on insulin signaling as shown by total Akt and its phosphorylation levels in inguinal white adipose tissue (K), brown adipose tissue (L), and gastrocnemius muscle (M). Each lane represents a pooled sample of three mice.

Further examination of adipose tissue histology revealed marked NAR effect on reducing adipocyte cell size in both the inguinal beige and visceral classical white adipose depots (Fig. 5A). More evident cytoplasmic staining in iWAT indicative of browning effect was observed in the beige depot. Immunoblot analysis of NAR-treated iWAT demonstrated markedly higher protein expression of RORα, BMAL1 and CRY2, indicating activation of the core loop of circadian clock gene regulation (Fig. 5B). Elevated levels of UCP-1, PGC-1α together with mitochondrial components SDHB and ATP5a provided further validation of the browning effects of NAR in (Fig. 5C). Similar to the effects observed in beige adipocytes, NAR treatment led to suppression of the lipogenic program together with reduced PPARγ level, although C/EBPβ was not altered as compared to controls (Fig. 5D). Lipolysis could be up-regulated in beige fat, as shown by marked inductions of ATGL and phosphorylation of HSL without altered HSL total protein level (Fig. 5E). RT-qPCR analysis further demonstrated the suppression of *Fasn* gene expression by NAR, suggesting a transcriptional repression mechanism in mediating this effect (Fig. 5F).

**Figure 5.**
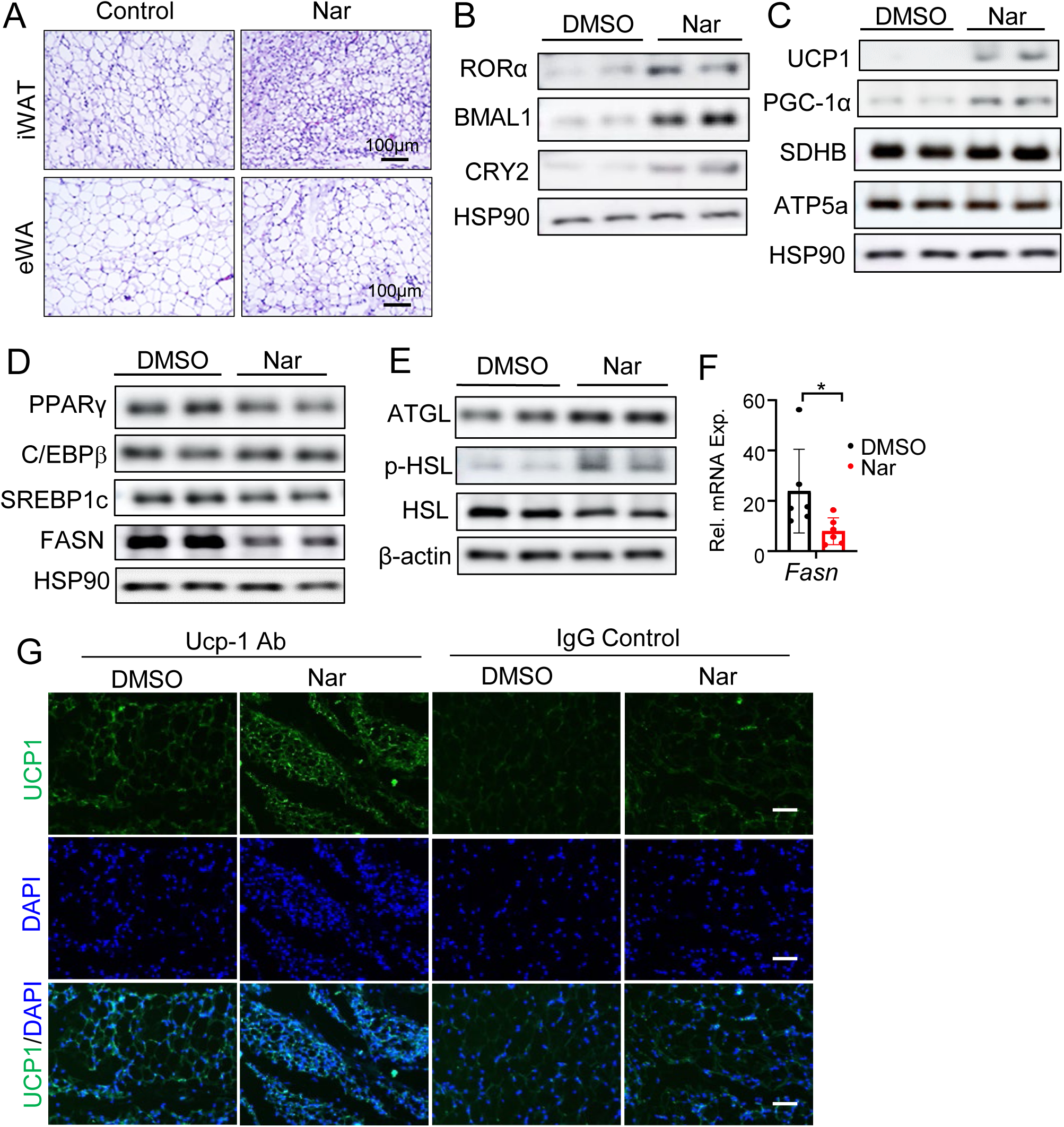
Naringenin treatment suppresses lipogenesis and induces browning of inguinal subcutaneous fat depot. (A) Representative H/E histology images of inguinal white adipose tissue (iWAT) and visceral epidydimal white adipose tissue (eWAT) in mice following Naringenin treatment for 7 days. (B-E) Immunoblot analysis of Naringenin effect on modulating clock regulators (B), thermogenic markers (C), lipogenic (D), and lipolytic proteins (E) in iWAT of mice following Naringenin treatment. (F) RT-qPCR analysis of fatty acid synthase (*Fasn*) in mice treated with Naringenin. N=6/group. *, **: P≤0.05 or 0.01 by Student’s *t* test. (G) Representative images of immunofluorescence staining of UCP-1 (green) in DMSO or Naringenin-treated inguinal subcutaneous fat. IgG was used as negative control for staining.

Consistent with the observed reduction of fat mass, in Nar-treated brown adipose tissue, there were similar reductions of lipid accumulation with stronger cytoplasmic staining (Fig. 6A). Interestingly, although NAR can induce RORα protein in brown fat similar as seen in white adipose tissues together with higher DBP protein indicative of increased core clock transcriptional output, it was not sufficient to induce Bmal1 level in BAT (Fig. 6B). Examination of lipogenic program in BAT revealed near complete suppression of FASN together with markedly attenuated ACC level, while both C/EBPβ and PPARγ were also significantly down-regulated (Fig. 6C). In contrast to beige adipose tissue, the protein expression of lipolytic enzymes, ATGL, HSL along with its phosphorylation in BAT were moderately reduced, suggesting NAR administration suppressed lipid remodeling while its inhibition of lipid accumulation was more dominant. Interestingly, UCP-1, PGC-1α and SDHB protein level were not significantly altered, suggesting that the browning effect of NAR is specific to beige fat depots without altering thermogenic regulation in the brown fat (Fig. 6D). In line with their reduced protein levels, analysis of *Srebp-1c* and *Fasn* mRNA indicated strong transcriptional repression of the lipogenic program by NAR (Fig. 6E).

**Figure 6.**
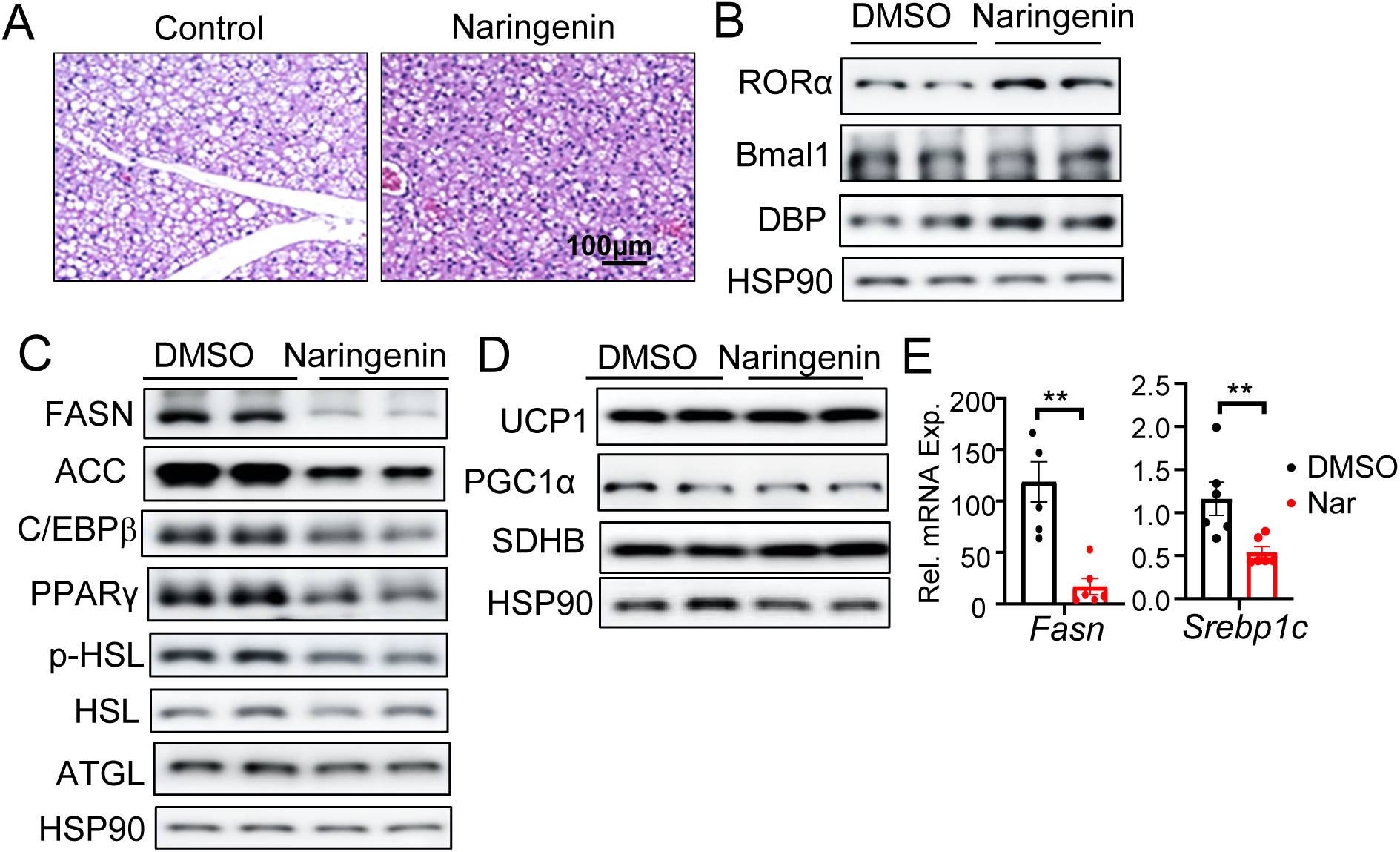
Effect of Naringenin on suppressing lipogenesis in brown adipose tissue. (A) Representative H/E histology images of brown adipose tissue (BAT) following 7 days of Naringenin treatment. Scale bar: 100 μm. (B-D) Immunoblot analysis of the effect of Naringenin on modulating clock regulators (B), lipogenic and lipolytic proteins (C), and thermogenic markers (D) in brown adipose tissue. (E) RT-qPCR analysis of lipogenic gene expression in mice treated with Naringenin. *, **: P≤0.05 or 0.01 by Student’s t test.

## Discussion

Despite the known beneficial metabolic effects of natural flavonoid molecules, their underlying mechanisms of action warrant detailed mechanistic investigations. Our current study identified a novel mechanism of action of NAR, an abundant citrus flavonoid, function as a RORα agonist to promote clock oscillation. We further demonstrate that NAR suppression of lipid storage in mature adipocytes is dependent on RORα, while its induction of mitochondrial activity led to adipocyte browning. Importantly, in vivo administration of NAR resulted in markedly reduced adipocyte lipogenesis with browning of beige of fat depot. Thus, the mechanisms of actions of NAR in adipocytes may collectively contribute to the observed anti-obesity and insulin-sensitizing efficacy of NAR in vivo. The discovery of the clock-modulatory activities of NAR implicates its potential applications for metabolic disease management, particularly as supplemental regimen in specific populations, such as shift workers, who are susceptible to circadian misalignment-induced metabolic consequences (Chaput et al., 2023; Luckhaupt, Cohen, Li, & Calvert, 2014).

Our investigation of NAR activity as a potential RORα-activating ligand is based on its shared flavonoid structural scaffold with Nobiletin, a known clock modulator that was identified to function as a RORα agonist (He et al., 2016). Naringenin displayed bona-fide RORα agonist activity in a RORE-driven luciferase reporter assay, albeit less potent than a synthetic ligand SR1078. Using RORα-null adipocytes, we demonstrated that the effect of Nar on inhibiting lipid store is RORα-dependent. RORα is a ligand-dependent nuclear receptor that activates gene transcription with diverse metabolic functions, although may also modulate gene regulation via specific interactions with other transcription factors (Jetten, 2009; Kim et al., 2017). Interestingly, despite the anti-obesity effects of NAR observed in previous studies, its clock-modulatory activity and the role in mediating distinct metabolic processes in adipocytes are not known. The identification of NAR as a RORα agonist reveals its specific target in the molecular clock feedback circuit. As a RORα ligand, transcriptional targets of RORα may mediate, at least in part, the beneficial effects of NAR and thus providing a tractable mechanism of action for NAR in metabolic regulation. Based on the current findings, specific RORα metabolic targets in mediating the in vivo action of NAR warrant further investigations. Nonetheless, given the multitude of metabolic benefits of NAR reported to date, including its robust anti-oxidant and anti-inflammatory properties (Alam et al., 2014; Mulvihill, Burke, & Huff, 2016), it is likely that RORα-mediated actions may only partly account for the observed NAR benefits in vivo.

Consistent with its ability to activate RORα and regulation of a direct RORE response element in Bmal1 promoter, NAR was able to induce Bmal1 protein expression together with up-regulation of its heterodimer partner CLOCK (Preitner et al., 2002). Interestingly, in addition to stimulate RORα activity as an agonist, NAR was able to induce RORα expression in adipocytes and in vivo, suggesting a potential positive regulatory loop involved in RORα gene transcription. RORα and the RORE-binding repressor Rev-erbα exert positive and negative modulations to drive Bmal1 oscillation, respectively (Guillaumond, Dardente, Giguere, & Cermakian, 2005). Bmal1 transcription activation by NAR modulation of RORα may result in secondary induction of RORα transcription, thereby forming a feed-forward mechanism in promoting clock oscillation via NAR modulation of RORα. Thus, RORα activation by NAR with inductions of Bmal1 as well as RORα itself may underlie its robust clock amplitude-enhancing activity. Further investigation of the precise regulatory dynamics of this regulatory loop, considered a secondary loop to provide redundancy of the core clock regulatory circuit, may reveal additional opportunities to target this mechanism for metabolically beneficial clock-enhancing interventions.

Combining in vitro testing of NAR in adipocytes with in vivo findings, our current study indicates that the actions of NAR in suppressing lipogenesis together with its ability to induce beige adipocyte browning may collectively contribute to the observed robust effects on reducing fat mass in mice. Key components of the molecular clock regulatory network exert distinct roles in modulating the development of specific types of adipocytes (Nam, Chatterjee, et al., 2015; Nam, Guo, et al., 2015; Nam et al., 2016; Xiong et al., 2021). NAR modulation of RORα is consistent with the known activity of RORα in blocking adipogenesis. A direct interaction of RORα with adipogenic factor C/EBPβ has been reported, leading to the inhibition of a key step in adipogenic induction of adipocyte progenitors (Ohoka et al., 2009). As a RORα agonist, NAR may inhibit adipogenesis by blocking C/EBPβ activity, although transcriptional effects of RORα, particularly its direct targets involved in lipid metabolism, may also mediate actions of NAR. The staggerer (*SG/SG*) mice, a RORα mutant, displays a lean phenotype, though various defects in this model precludes the direct assessment of RORα function in adipocytes (Lau et al., 2008; Lau, Nixon, Parton, & Muscat, 2004). In hepatocyte, RORα can suppress the lipogenic program by interfering with PPARγ activity through interaction with HDAC3, with negative regulations of *SREBP-1c* and *Fasn* (Kim et al., 2017). As a result, hepatic RORα deficiency resulted in severe hepatic steatosis with excessive fat accumulation, in line with NAR inhibition of lipogenic gene program in mature adipocytes. In addition, the attenuated effect of NAR on reducing lipids synthesis in mature adipocytes with RORα deficiency further supports this RORα-dependent mechanism. Potential overlap between targets of RORα in lipogenic regulation in hepatocyte with adipocytes remains an intriguing possibility.

Our findings revealed that NAR was able to induce significant loss of adipose tissue within a short period of one week administration. NAR exerts robust effect on adipose tissue remodeling, suppressing lipid synthesis while promoting lipolytic activation and browning activity in beige adipocytes. These actions could be synergistic in reducing lipid storage while enhancing fatty acid oxidation through mitochondrial activity. Interestingly, despite the up-regulation of lipolytic enzymes in adipose tissues, serum free fatty acid level in NAR-treated mice remains lower than that of controls, suggesting potential enhanced oxidation by skeletal muscle. Nobiletin enhanced mitochondrial activity in muscle that contributes to its anti-obesity effect (Nohara et al., 2019), and NAR displays similar effect on inducing browning of subcutaneous depots. Given the reported pleotropic actions of flavonoid compounds, additional mechanisms may also apply. We found similarly enhanced insulin signaling by NAR treatment in both adipose tissue and skeletal muscle, suggesting, at least in part, overlapping mechanisms underlying the metabolic regulations. Additional analysis in the skeletal muscle confirmed induction of the clock circuit and Glut4 protein level (data not shown). The increase in Akt phosphorylation by NAR in adipose tissue and skeletal muscle suggests insulin-sensitizing actions in distinct peripheral tissues, potentially in the liver as well, may contribute in aggregate to the observed improvement in systemic glucose homeostasis. Whether and to what extent these regulations are mediated by NAR activation of RORα in vivo remains to be determined.

Our current findings are, in large part, in agreement with the reported benefits of NAR with lipid-lowering, anti-atherosclerosis and anti-obesity effects (Ke et al., 2015; T. Pan et al., 2024; Rebello et al., 2019; Ucar Bas et al., 2024). This study provides a novel mechanistic connection of NAR as a RORα-activating molecule that exerts clock-modulatory actions in suppressing fat storage and enhancing lipid mobilization. While focused on its actions in adipocytes, this specific mechanism in mediating NAR effects may apply to metabolic processes in liver or the vasculature that contributes to its robust lipid-lowering efficacy in vivo (Mulvihill et al., 2016). Several recent studies reported the activity of NAR in inducing browning of beige adipocytes, albeit largely limited to cellular models that failed to interrogate the underlying mechanisms (Rebello et al., 2019; Ucar Bas et al., 2024). Our findings of NAR as a direct RORα-activating natural ligand thus provides novel mechanistic insights into this metabolic beneficial effect on thermogenic induction. Given the known multifaceted metabolic actions of NAR and RORα function, these studies warrant further investigation of the extent to which RORα activation mediates these actions of NAR in thermogenic fat depots in vivo. A better understanding of RORα-mediated mechanisms involved may also add to the current repertoire of the known metabolic activities of NAR. The direct actions of NAR on inhibiting lipid remodeling in adipocytes with its anti-obesity efficacy in vivo suggests this natural compound may have potential for therapeutic intervention or prevention of obesity and its associated metabolic complications, particularly considering its established safety profile. With the increasing prevalence of circadian disruption in our society (A. Pan, Schernhammer, Sun, & Hu, 2011; Roenneberg, Allebrandt, Merrow, & Vetter, 2012), it is also possible that the clock-modulatory activities of NAR could be leveraged to re-enforce clock function for long-term interventions for metabolic diseases associated with circadian misalignment.

## Acknowledgements

We greatly appreciate the kind gift from Dr. Dan Littman at NYU Langone School of Medicine by providing us with the floxed RORα mouse line. We thank the City of Hope Shared Resources Animal Phenotyping for carrying out metabolic phenotyping analysis. KM is a faculty member supported by the NCI-designated Comprehensive Cancer Center at the City of Hope National Cancer Center. This project was supported by National Institute of Health grants R01DK112794, R56AG080294, an Innovative Award from Arthur Riggs Diabetes and Metabolism Research Institute to KM. VY is supported by National Institute of Health grants R01DK097160 and R01DK128972, and AP is supported in part by National Institute of Health grant R01DK128972. These funders had no role in study design, data collection and analysis, decision to publish, or preparation of the manuscript.

## Authorship Statement

XX, JP and TK: data curation and investigation, formal analysis, manuscript editing; ZS, AP and VY: data curation and manuscript editing; KM: formal analysis, project administration, manuscript writing and editing, and funding acquisition.

