## Supplementary material for "Clock Modulation by Naringenin via RORα Suppresses Lipogenesis and Promotes Adipose Tissue Browning": Suppl. tables

**Supplemental Table 1. Primary antibodies list.**

| Antibody | Source | Cat# | Dilution |
|---|---|---|---|
| BMAL1 | Santa Cruz | SC-365645 | 1:1000 |
| RORα | Proteintech | 82930-1-RR | 1:1000 |
| DBP | Proteintech | 12662-1-AP | 1:1000 |
| CRY2 | Invitrogen | PA5-79073 | 1:1000 |
| AKT | Cell Signaling | 4691S | 1:1000 |
| p-AKT(S473) | Cell Signaling | 4060S | 1:1000 |
| HSL | Cell Signaling | 4107S | 1:1000 |
| p-HSL(S563) | Cell Signaling | 4139S | 1:1000 |
| ATGL | Cell Signaling | 2439S | 1:1000 |
| UCP1 | Cell Signaling | 72298S | 1:1000 |
| PGC1a | Sigma | ST1204 | 1:1000 |
| ACC | Cell Signaling | 3676S | 1:1000 |
| ATP5a1 | Proteintech | 14676-1-AP | 1:1000 |
| SDHB | Proteintech | 10620-1-AP | 1:1000 |
| SREBP1 | Santa Cruz | SC-13551 | 1:1000 |
| C/EBPβ | Santa Cruz | SC-7962 | 1:1000 |
| PPARγ | Santa Cruz | SC-7273 | 1:1000 |
| FASN | Santa Cruz | SC-48357 | 1:1000 |
| FABP4 | Santa Cruz | SC-271529 | 1:1000 |
| HSP90 | Cell Signaling | 4874S | 1:3000 |
| β-Actin | Proteintech | 66009-1-Ig | 1:3000 |

**Supplemental Table 2. Primer sequence for qPCR analysis.**

| **Genes** | | **Sequences** |
|---|---|---|
| Fasn | Forward | GTGAGTCTATCCTGCGCTCC |
| | Reverse | CCCAAGGAGTGCCCAATGAT |
| Srebp-1c | Forward | ACCACGGAGCCATGGATTG |
| | Reverse | CTGTGTCCCCTGTCTCACC |
| 36B4 | Forward | CGCTTTCTGGAGGGTGTCCGC |
| | Reverse | TGCCAGGACGCGCTTGTACC |
